# Novel “*Candidatus* Endobugula” hosts and distinct microbiomes in the bryozoans from the White Sea

**DOI:** 10.64898/2026.09.15.751706

**Authors:** Anna M. Dukat, Egor S. Trubnikov, Evgeniy A. Bogdanov, Olga N. Kotenko, Andrey E. Vishnyakov, Mikhail S. Gelfand, Andrew N. Ostrovsky

## Abstract

Marine bryozoans harbor diverse microbial communities, but not much is known about specificity of bacterial symbionts towards different hosts. We identified potential symbionts and described overall microbial communities of three marine bryozoans, *Aquiloniella scabra, Rhamphostomella bilaminata* (Gymnolaemata: Cheilostomatida), and *Patinella verrucaria* (Stenolaemata: Cyclostomatida), revealing that each species maintains a distinct microbial assemblage that differs both from the surrounding environmental microbiome and from other hosts. Previously, the bryostatin-producing gammaproteobacterial genus “*Candidatus* Endobugula” was known from only two bryozoan species in the family Bugulidae. Here, we report its first detection in two more cheilostome bryozoans, *A. scabra* and *R. bilaminata*, expanding its known host range to two more families.

## Introduction

Marine invertebrates are known for hosting complex and diverse microbial communities that can provide metabolic capabilities that their hosts lack. This diversity and specificity of the invertebrates’ microbiome make it difficult to identify important symbionts and to understand how these communities are structured [1].

Some symbioses are dominated by a single endosymbiont that is largely responsible for the host’s wellbeing. The best-studied cases are in insects: mutualists such as *Buchnera* [2] and *Wigglesworthia* [3] are obligate, bacteriocyte-housed bacteria that synthesize essential nutrients limited in the host’s diet, whereas parasitic symbionts such as *Wolbachia* propagate not by feeding the host but by manipulate its reproduction to favor infected females that transmit them [4]. In all these instances, symbionts are transmitted by the new host generation using vertical transfer.

The same single-partner arrangement is found in marine environments, but with a crucial difference: vertical inheritance of the symbiont is not guaranteed. The Hawaiian bobtail squid maintains an exclusive association with one bacterial species, *Vibrio fischeri*, housed in a specialized light organ [5], and vestimentiferan tube worms are also generally mono-symbiotic with sulfur-oxidizing chemoautotrophs living in a trophosome [6]. In both cases, however, a tight and specific host–symbiont relationship is rebuilt each generation from the environment (environmental transfer) rather than inherited by vertical transfer [7].

A mixed model occurs in some corals, where a community of the algal symbionts from the genus *Symbiodinium* is partly inherited vertically and partly acquired from the environment [8]. This holds even when a host simultaneously carries several symbionts. Gutless oligochaete worms host a consortium of distinct chemosynthetic bacteria [9]. The whole consortium is transmitted from parent to offspring, but with fidelity that varies member by member: dominant, nutritionally essential sulfur-oxidizers are always inherited, while lower-abundance partners are transmitted only loosely or re-acquired anew [10].

At the other end of this gradient are hosts associated with diverse microbial communities rather than one or a few dominant partners. Corals harbor rich bacterial assemblages whose composition shifts with season and dormancy [11], summer-winter cycles [12], geographic location [13], and can be further modulated by their parasites, such as gall-inducing copepods [14]. Sponges show the same pattern: a large, structured, and convergently evolved microbiome shared across host species worldwide [15].

These differences between the symbiont inheritance/transmission in aquatic and terrestrial habitats are shaped by the habitat itself: across more than 500 documented bacteria-eukaryote symbioses, strictly vertical inheritance is abundant among terrestrial hosts and scarce among aquatic ones, where the environmental and mixed modes prevail [16]. A likely reason is availability. In water, symbionts persist as free-living populations that each new host generation can acquire anew: vent tubeworm larvae hatch symbiont-free and must be colonized from the environment through the epidermis [17], and deep-sea mussels endocytose free-living sulfur-oxidizers at the gill surface [18].

Most known cases from another filter-feeding group, phylum Bryozoa, are persistent mono-symbioses [19–21]. The best described symbiosis is the association between cheilostome bryozoan *Bugula neritina* and uncultured gammaproteobacterium “*Candidatus* Endobugula sertula”, known for bryostatin production [21]. This partnership was described as inherited rather than re-acquired: the symbiont population is acquired by the host’s larvae from the parent during brooding, carried in their pallial sinus and being passed to the daughter colony during larval metamorphosis [22, 23]. However, unlike other obligate, vertically transmitted, host-restricted symbionts, “*Ca*. Endobugula sertula” shows no genome reduction [24], possibly indicating recent symbiosis or persistence of the symbiont in the environment. A recent study questioned the widely accepted conclusion on the vertical transfer in some bryozoans [25]. This is in accord with the fact that “*Ca*. Endobugula sertula” was detected in the environmental control of sea water near *B. neretina* colonies, indicating possible acquisition of the symbiont from the environment [20].

Bryostatins defend Bugula’s larvae from predation, so the main demonstrated function of this microbial symbiosis is chemical defense [22]. Bryostatins are macrolactones that act mainly by modulating protein kinase C and show antitumor, immunomodulatory, and anti-inflammatory activity [26]. Prior to this study, only two bryozoan species, *B. neritina* and *Bugulina simplex* (Bugulidae) were known to host bryostatin-producing symbionts, “*Ca*. Endobugula sertula” and “*Ca*. Endobugula glebosa” respectively [23].

A second well-studied case is the association between bryozoans of the genus *Watersipora* and “*Candidatus* Endowatersipora”. The symbionts, first described as mollicutes in *W. arcuata*, were reassigned based on the 16S rRNA gene sequencing to *Alphaproteobacteria*, and named “*Ca*. Endowatersipora palomitas” and “*Ca*. Endowatersipora rubus” for the symbionts of cheilostomes *W. subtorquata* and *W. arcuata* respectively [27, 28]. By analogy with “*Ca*. Endobugula”, it has been suggested that these bacteria may contribute to the chemical defense of host larvae, though this remains untested [27]. Bioassay-guided fractionation of *W. subtorquata* yielded bryoanthrathiophene along with two related anthraquinone-type compounds, of which bryoanthrathiophene showed the strongest antiangiogenic activity in an endothelial-cell proliferation assay [29]. Whether these metabolites are of bacterial origin, and whether they serve a defensive role in the host, has not been established.

Not all bryozoans follow this pattern. The Arctic cheilostome bryozoan *Tricellaria ternata* harbours a complex bacterial community, distinct from those of the surrounding water and sediment, but with no sign of any persistent mono-symbiosis [30]. The freshwater bryozoan *Cristatella mucedo* likewise lacks a dominant single symbiont, and its community overlaps considerably with those of the surrounding water and sediment; however, bacterial isolates have proven secondary-metabolite biosynthetic potential, and metabolomic analysis of mono- and co-cultures recovered both known and potentially novel compounds [31].

Here, we provide the first description of microbiomes of three marine bryozoans from two distant clades, *Aquiloniella scabra, Rhamphostomella bilaminata* (class Gymnolaemata: order Cheilostomatida), and *Patinella verrucaria* (class Stenolaemata: order Cyclostomatida), for the first time. We also identify their potential symbionts previously recorded by the microscopic methods [32, 33] and discuss their origin.

## Materials and Methods

### Site description and sampling

All samples were collected in the first half of June 2025 in the vicinity of the Educational and Research Station “Belomorskaia” (White Sea, Kandalaksha Gulf, Chupa Inlet). Detailed collection dates, sampling techniques, site coordinates, and substrate type for each sample are provided in Table 1.

**Table 1.**
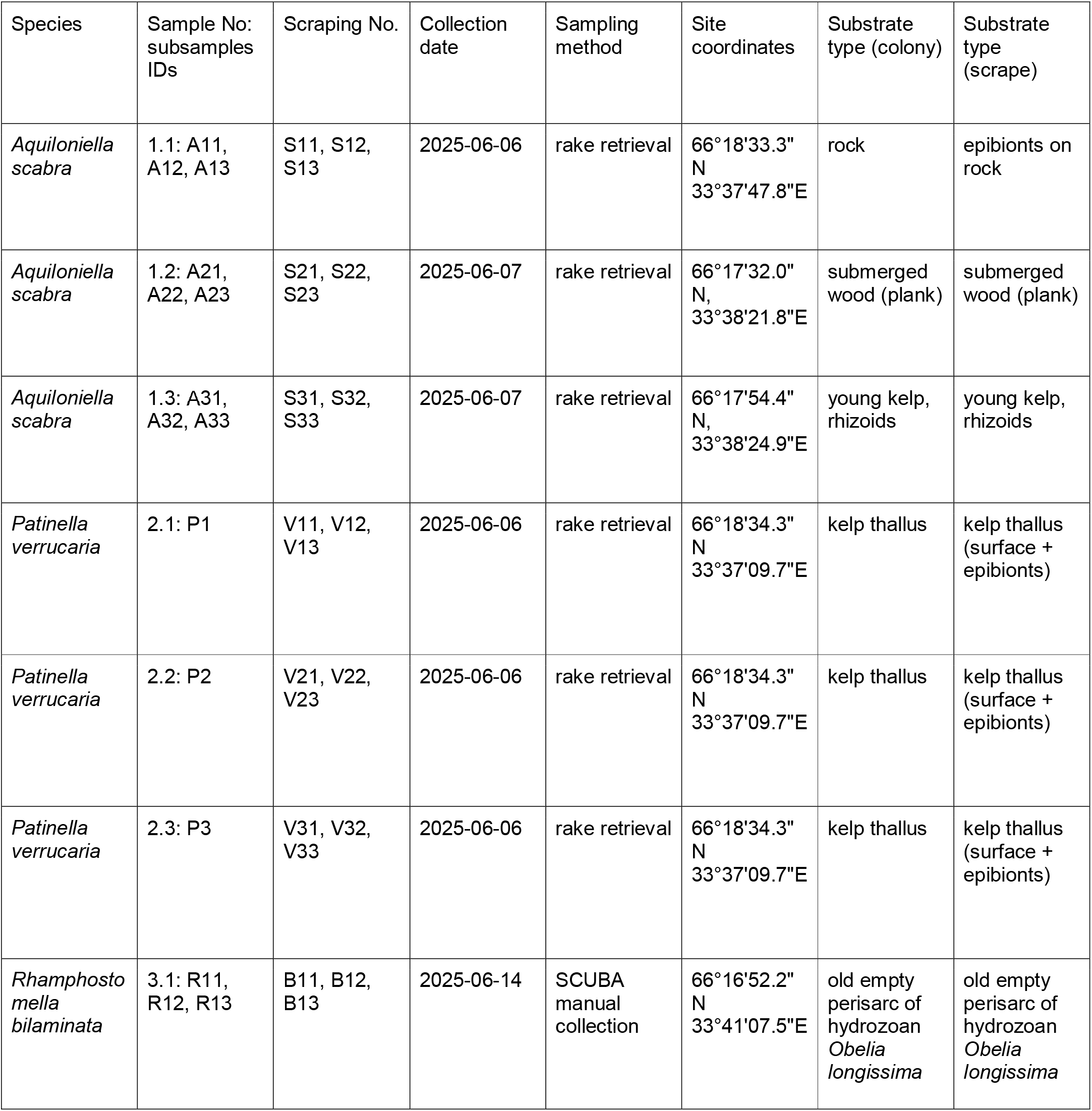
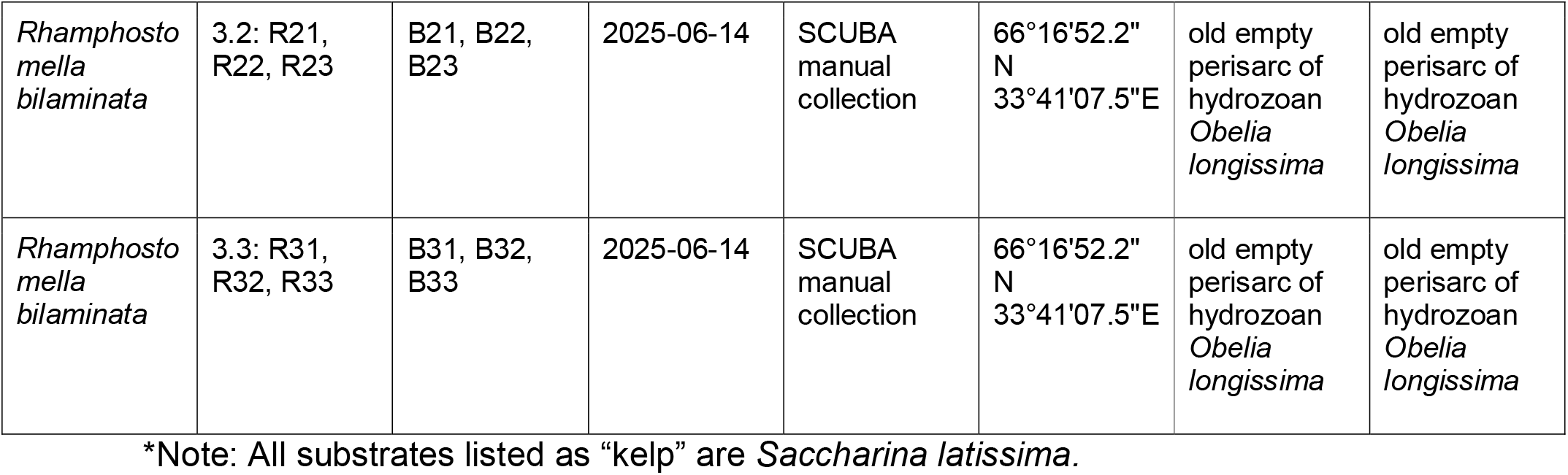
Description of the sampling sites.

Three bryozoan species were chosen: erect cheilostome *Aquiloniella scabra* (van Beneden, 1848), encrusting cheilostome *Rhamphostomella bilaminata* (Hincks, 1877), and knob-like cyclostome *Patinella verrucaria* (Linnaeus, 1758). Three specimens of each species were collected; for each specimen, substrate surface scraping was performed using sterile swabs in three replicates: the first scraping was taken directly adjacent to the colony attachment site, the second at a distance, and the third further away (Table 1). All samples were placed in sterile filtered buffer containing 10 mM Tris, 0.1 M EDTA, and 0.5% SDS (pH 8.0) and stored at 0-6 °C. Samples were processed further two weeks following fixation. During sample preparation, three fragments were cut from each *A. scabra* and *R. bilaminata* colony sample. Compact colonies of *P. verrucaria* were not sectioned. Live colonies were photographed using a DS-Fi3 camera attached to a Nikon SMZ745T stereomicroscope.

### DNA extraction and sequencing

DNA was extracted using the QIAamp Fast DNA Tissue Kit (Qiagen, Germany) from 25 mg of each sample. Samples were homogenized for 5 min using vertical homogenizer TissueLyser LT (Qiagen, Germany). Following sample disruption, DNA extraction was performed according to the manufacturer’s instructions. DNA concentrations were measured using the Qubit fluorometer (Thermo Fisher Scientific, USA) with the dsDNA HS assay kit (Thermo Fisher Scientific, USA). The extracted DNA was stored at −20 °C until the start of the amplicon library preparation. To account for the possible contamination, negative control with sterile water was used for the amplification.

The V4 variable region of the 16S rRNA gene was amplified using the universal primers 515F 5′-GTGCCAGCMGCCGCGGTAA-3′ and 806R 5′-GGACTACHVGGGTWTCTAAT-3′ [34]. To avoid the adaptor ligation step, Illumina adaptors 1 and 2 were added to the primer sequences. Phusion 2X Master Mix with the HS buffer (New England Biolabs, USA) was used for amplification. The DNA was denatured (95 °C; 2 min), followed by 28 cycles: denaturation (95 °C; 25 s), annealing (58 °C; 30 s), extension (72 °C; 40 s), then final extension (72 °C; 3 min) and holding (4 °C). Amplicons were then purified with the AMPure XP beads (1:0.9 V:V, Beckman Coulter), and their concentration was measured on the Qubit fluorometer. To control possible contamination, negative PCR control was made with sterile water. The Nextera XT index kits (Illumina, USA) were then used to index the amplicons. The resulting libraries were cleaned using the AMPure XP beads, quantified, and sequenced in the Skoltech Genomics Core Facility on Illumina MiSeq (Illumina, USA) as 250 PE using the MiSeq reagent kit V2 (500 cycles).

### Sequencing data analysis

Demultiplexed paired-end raw reads were processed using QIIME2 (version 2026.1) [35]. Primer sequences were trimmed from the reads using the q2-cutadapt plugin [36]. Trimming was performed with a maximum error rate of 0.2, allowing matches to both read and adapter wildcards, and all untrimmed reads were discarded. Following adapter and primer removal, per-base quality profiles of the trimmed sequences were assessed using FastQC [37] and aggregated using MultiQC [38] to confirm read integrity prior to denoising. Denoising, chimera checking, and paired-end read merging were performed using the DADA2 plugin [39]. Taxonomy was assigned to the ASVs (amplicon sequence variants) using the q2-feature-classifier plugin [40] with a scikit-learn naive Bayes classifier trained on the SILVA database (v138) [41]. Multiple sequence alignment of the representative ASV sequences was generated using MAFFT [42], and a phylogenetic tree was constructed using FastTree [43], both as implemented via the q2-phylogeny plugin. Finally, the ASV feature table, representative sequences, taxonomy assignments, and rooted phylogenetic tree were exported from QIIME2 for downstream statistical analyses.

### Statistical data analysis

All subsequent analyses were performed in R version 4.6.0 (2026-04-24). QIIME2 artifacts (feature table, taxonomy, rooted phylogenetic tree) were imported into R using the qiime2R package. ASV table, taxonomy assignments, and metadata were combined into a phyloseq object using the phyloseq package (version 1.44.0) [44]. PCR controls were identified, and contaminant ASVs were removed using the prevalence method implemented in the decontam package (version 1.20.0) [45] with a prevalence threshold of 0.5. After decontamination, ASVs present in fewer than two samples (singletons) were filtered out. The clean feature table was then transformed to relative abundances for visualization and ordination.

Alpha-diversity was assessed after rarefying the clean ASV table to the minimum sequencing depth (minimum library size was 20337 reads). To account for variation in library sizes among samples and characterize random variation introduced by rarefying, we implemented a repeated rarefaction approach [46]. We repeatedly rarefied each sample 100 times across 20 depth points ranging from 10% of the sample’s library size to its total library size. The normalized library size was set to the minimum library size among all samples after filtering, ensuring equal sequencing depth for alpha-diversity comparisons.

Beta-diversity was explored using non-metric multidimensional scaling (NMDS) on Bray-Curtis distances calculated from relative abundance data (log-transformed with log1p), using the metaMDS function from the vegan package (version 2.6-4) [47]. PERMANOVA (adonis2 from vegan) was performed to test for differences between sample types (host vs. control) within each species, with 999 permutations. Pairwise distances were computed for both Bray-Curtis (relative abundance) and Jaccard (presence/absence) metrics. For each host species, distances were categorized as: within the same species, between different species, and sample vs. environmental control.

Differential abundance analysis was performed using MaAsLin2 (version 1.16.0) [48]. For each host species individually, the ASV counts (filtered to prevalence ≥2 samples) were compared between host samples and environmental controls, with sample type as a fixed effect and environmental controls set as the reference level. Significant ASVs were identified at a false discovery rate (FDR) p-value < 0.05. Chloroplasts, mitochondria, and ASVs not assigned to a class level were removed.

Phylogenetic reconstruction of the “*Ca*. Endobugula” 16S rRNA V4 region was performed using the Maximum Likelihood method implemented in MEGA12 [49]. Nodal support was evaluated with 1,000 bootstrap replicates.

### Transmission electron microscopy

For light and transmission electron microscopy (TEM), individual colony fragments of *Aquiloniella scabra* and *Rhamphostomella bilaminata* were fixed in 2.5% glutaraldehyde (buffered in 0.1 M Na-cacodylate with 10.26% sucrose, pH 7.4) for 1 hour, washed three times in buffer each lasting 15 minutes, and postfixed in 1% osmium tetroxide (OsO4) during 1 hour followed by three rinses in distilled water, each lasting 20 minutes. Fragments were then decalcified in 8.5% EDTA solution from a few hours to one day following by washing in distilled water. The dehydration process involved an ethanol series (30-50-70-80-90-100%) and acetone, after which they were embedded in epoxy resin type TAAB 812. Semithin sections (1.0 µm thick) were made using a ultramicrotome Leica EM UC7 (Leica Microsystems, Wetzlar, Germany), stained by the Richardson [50] and Humphrey & Pitman [51] techniques and examined with a light microscope Leica DM 2500. Ultrathin sections (70 nm thick) were made using a Leica EM UC7 microtome. The sections were collected on copper grids and contrasted in uranyl acetate and lead citrate being further examined using JEOL JEM-1400 and JEOL JEM-2100HC (JEOL Ltd., Japan) transmission electron microscopes and photographed with digital CCD cameras.

## Results

### Microscopic observation of bacteria associated with bryozoans

In the colonies of cheilostomes *A. scabra* and *R. bilaminata*, bacteria were found on the external zooidal walls and the wall of the vestibulum (a small chamber below the zooidal opening through which the tentacle crown is protruded). Gram-negative bacteria on the colony surface were rather diverse morphologically being from round and oval to elongate in shape. They were randomly distributed and never formed groups. In contrast, groups of oval bacteria of the same morphotype were recorded in both studied bryozoans in the folds of the vestibular wall in the feeding autozooids (Fig. 1A). In addition, in *A. scabra*, bacteria were also found inside the zooidal visceral cavity, occurring in the specialized organs, “bacterial bodies” (Fig.1B) [33]. Each such body is an oval capsule with 2-3-layered cellular wall surrounding its internal cavity with bacteria.

**Figure 1.**
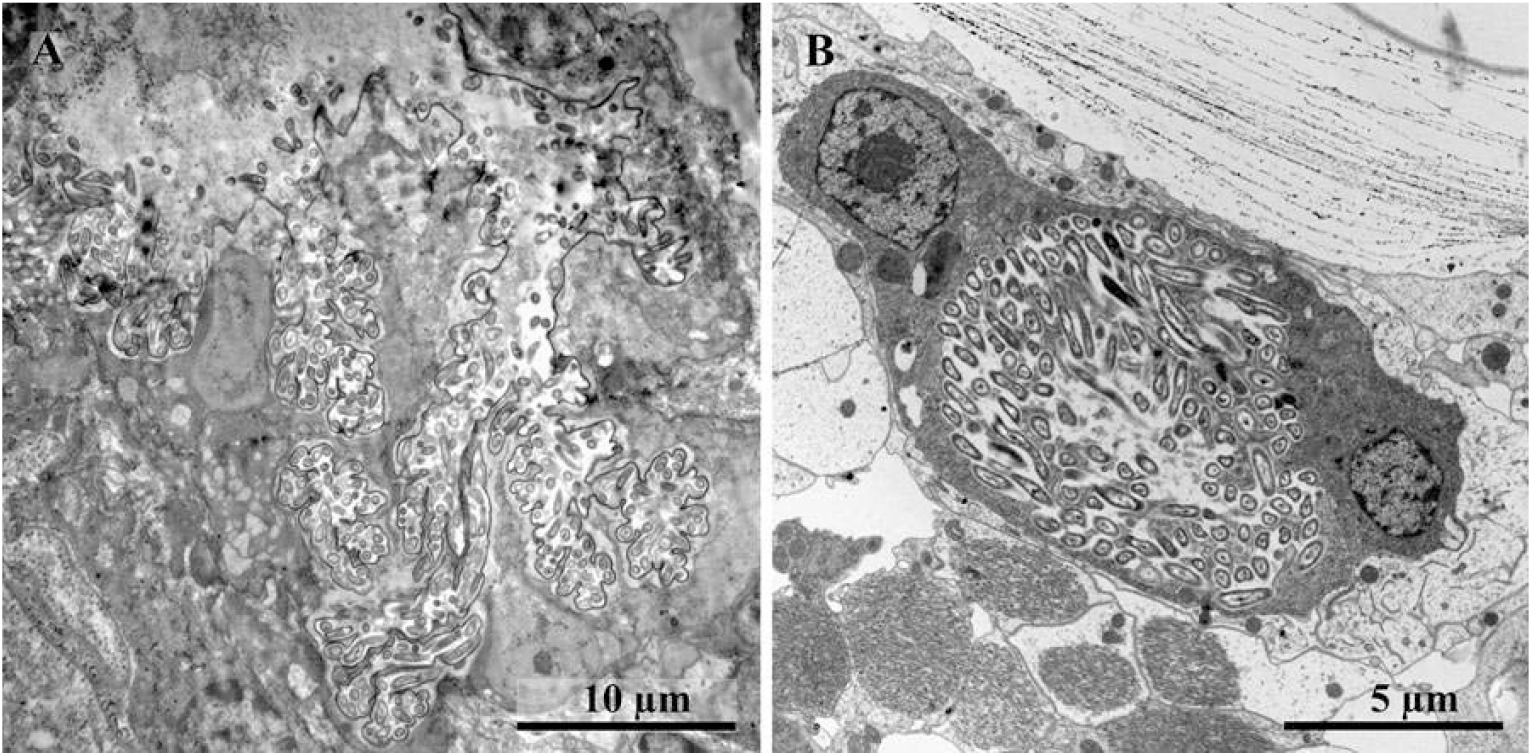
Bacteria associated with bryozoan autozooids (TEM). (A) *Rhamphostomella bilaminata*, vestibular folds filled with bacteria. (B) *Aquiloniella scabra*, intrazooidal bacteria inside the bacterial body.

In the cyclostome bryozoan *P. verrucaria*, intracellular bacteria were recorded inside tissues and individual cells (amoebocytes) [32].

### Microbiome composition of host bryozoans

The phylum composition was similar across all samples, including both environmental control and hosts, with predominant *Bacteroidota, Cyanobacteriota, Planctomycetota*, and *Pseudomonadota*.

At the genus level, major differences were observed between each bryozoan host and their respective environmental controls (Fig. 2). Chloroplast sequences were detected in high abundance across all bryozoan species, presumably reflecting their algal diet. In *A. scabra*, “*Ca*. Endobugula” was the most abundant genus relative to control. In *P. verrucaria*, the two most abundant genera relative to the control were *Yoonia-Loktanella* and *Sulfitobacter*. In *R. bilaminata*, top abundant taxa relative to control could not be characterized to the genus level.

**Figure 2.**
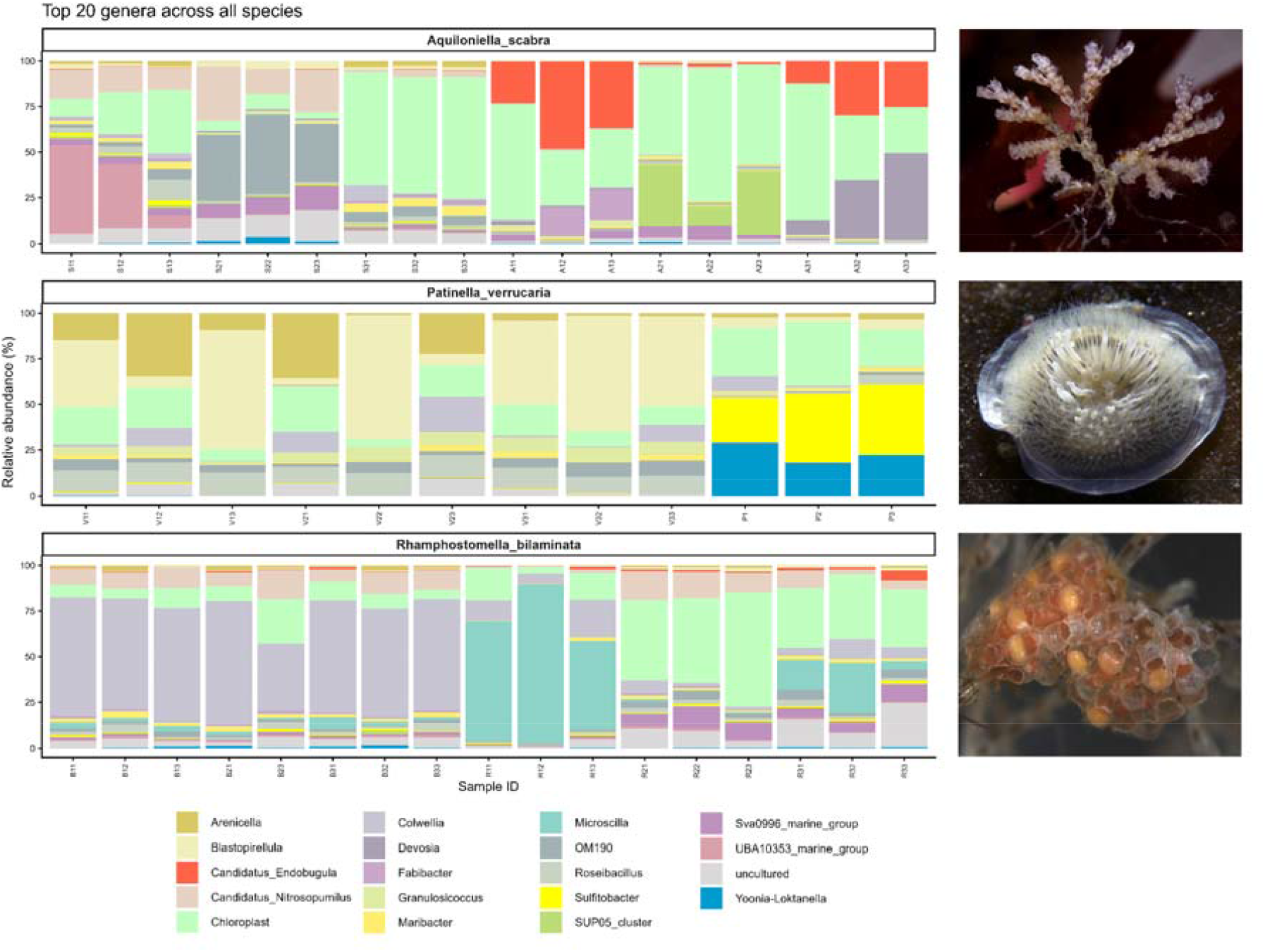
Taxonomic composition of the microbial communities of three studied bryozoan species at the genus level.

### Identification of “*Ca*. Endobugula” ASVs

Eleven ASVs assigned to the genus “*Ca*. Endobugula” were detected across the dataset. Two ASVs were consistently detected in two different bryozoan hosts; the V4 region of 16S rRNA sequences of “*Ca*. Endobugula” from of *A. scabra* (ASV_ASca, mean relative abundance of 14.6% across host replicates) and *R. bilaminata* (ASV_RBil, mean abundance of 0.5% across host replicates). In contrast, the remaining nine ASVs were rare and sporadic (Supplementary Table 1).

These two ASVs formed a moderately supported sister group in the phylogenetic tree (bootstrap support: 85.4%). This clade shared a deeper node with the reference sequences “*Ca*. Endobugula glebosa” (AY532642; terminal branch length: 0.00473) and “*Ca*. Endobugula sertula” (AF006606; terminal branch length: 0.03954), each of which branched independently from that node (Fig. 3). Despite close similarity of the V4 sequences, the abundance data revealed a strong, although uneven host-specific distribution: ASV_ASca was exclusively associated with *A. scabra*, whereas ASV_RBil was exclusively associated with *R. bilaminata*, neither ASV being detected in the other host or its environmental control, although they were seen in control samples of the respective hosts (Supplementary Fig. 1, Supplementary Table 1).

**Figure 3.**
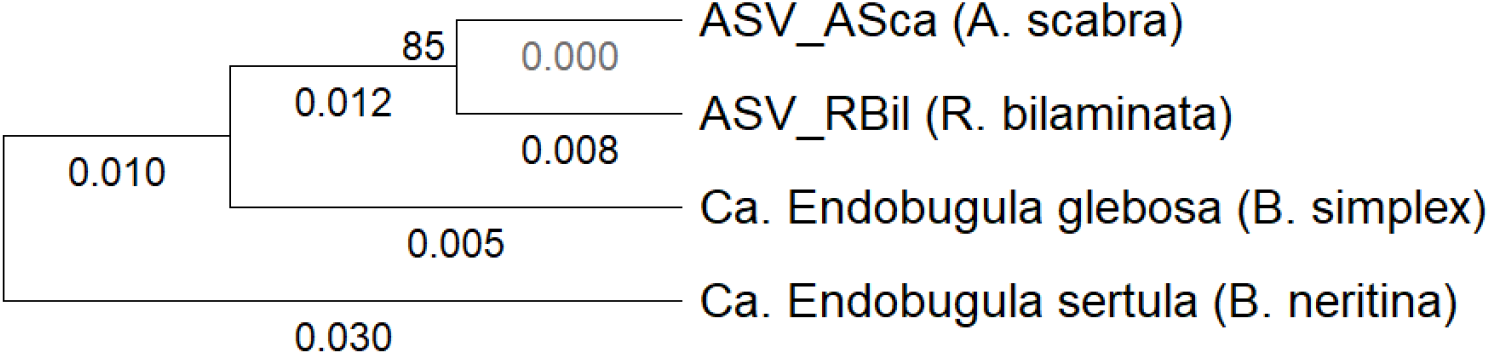
“*Ca*. Endobugula” maximum likelihood phylogenetic tree based on the V4 region of 16S rRNA.

### Alpha-diversity of bryozoan-associated microbiomes

Alpha-diversity was assessed to determine variation in microbiome richness, evenness, and taxonomy diversity between bryozoan samples and their respective environmental controls, using Observed taxa, Chao1, Shannon, and Faith’s Phylogenetic Diversity (PD) indices. To account for variation in the library sizes among samples, we performed repeated rarefaction and generated rarefaction curves for the alpha-diversity metrics. These curves indicated that the diversity estimates approached saturation at the selected sequencing depth (Supplementary Figure 2).

For *A. scabra*, all four alpha-diversity metrics were significantly lower in bryozoan samples than in environmental controls, whereas no statistically significant differences were detected after multiple-test correction for any of four alpha-diversity metrics for *P. verrucaria* and *R. bilaminata* (Supplementary Figure 3, Supplementary table Table 2).

### Beta-diversity reveals microbiome differences between hosts and environment

The NMDS ordination revealed distinct clustering, separating host samples from their respective controls while also discriminating among the host species (Fig. 4). The community structure was analyzed with PERMANOVA (centroid-based compositional shifts) to assess the separation and internal clustering of each species-associated assemblage relative to environmental controls. All PERMANOVA *p*-values were Bonferroni-adjusted for multiple comparisons. Beta-diversity was assessed using the Bray-Curtis (abundance-weighted) and Jaccard (presence/absence) dissimilarities.

**Figure 4.**
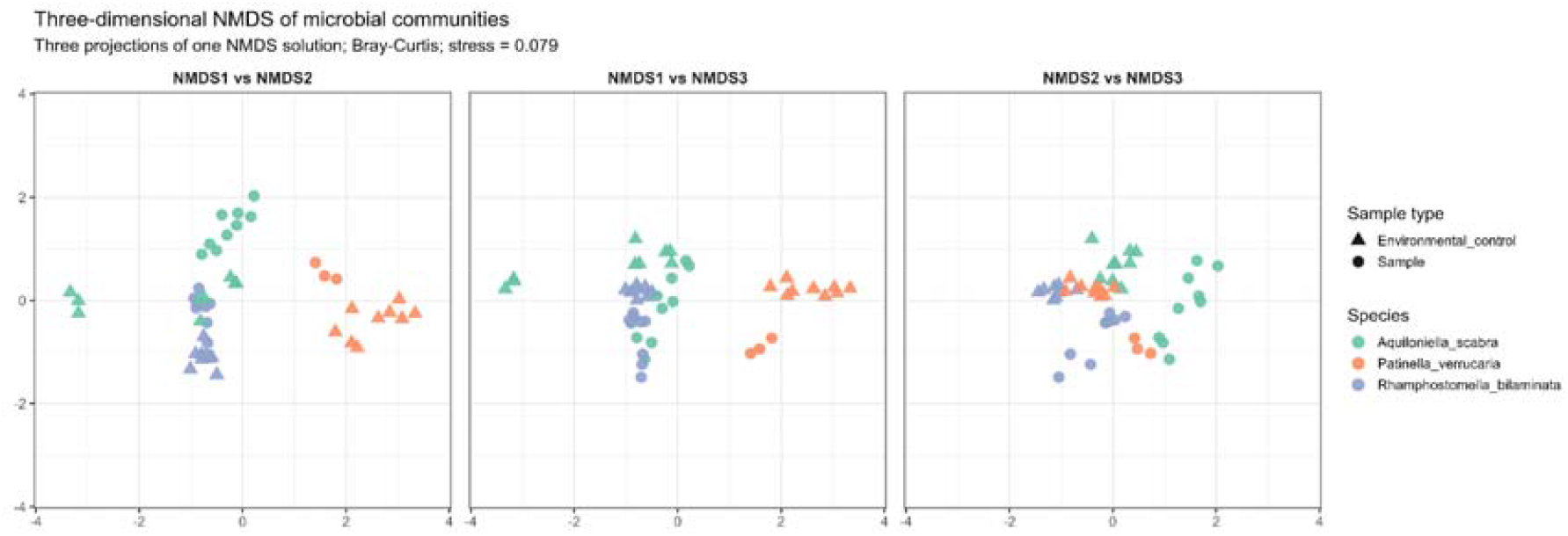
Three-dimensional Non-metric Multidimensional Scaling (NMDS) plot.

PERMANOVA revealed significant differences between host-associated microbial communities and their respective environmental controls for all three species (*A. scabra:* R^2^ = 0.210, Bonferroni-adjusted p = 0.003; *R. bilaminata*: R^2^ = 0.326, Bonferroni-adjusted p = 0.003; *P. verrucaria*: R^2^ = 0.429, Bonferroni-adjusted p = 0.021). Consistent with these results, pairwise Bray–Curtis and Jaccard distances were lower among samples from the same host species than between different host species or between hosts and environmental controls, indicating greater within-host community similarity. Medians of the Bray⍰Curtis distances within host species were consistently lower than the within⍰control distances (*A. scabra*: 0.749 for the sample and 0.919 for the control; *P. verrucaria*: 0.382 for the sample and 0.919 for the control; *R. bilaminata*: 0.620 for the sample and 0.919 for the control), suggesting that host microbiomes are more similar within species than the surrounding environmental microbiomes are. These patterns were consistent across both Bray-Curtis and Jaccard metrics (Supplementary Fig. 4).

### Differentially abundant taxa in bryozoan hosts

The number of differentially abundant ASVs (*p*-value < 0.05) across three host species included 5 enriched in host and 19 depleted ASVs in *A. scabra*, 62 enriched in host and 26 depleted ASVs in *P. verrucaria*, and 12 enriched in host and 32 depleted ASVs in *R. bilaminata*. Differential abundance analysis for *A. scabra* and *R. bilaminata* revealed a general pattern of depletion: more ASVs were significantly depleted in the host than were enriched in it. This suggests that host selection acts primarily as a filtering mechanism that excludes a broad range of microorganisms from the environment. However, *P. verrucaria* is different with more enriched than depleted ASVs, indicating that it either possesses weaker environmental filtering, or actively recruits a broader set of symbionts, or both. This species also harbors the highest total number of differentially abundant ASVs, suggesting a more variable and complex microbiome. Chloroplasts, mitochondria, and ASVs not assigned to a class level, were removed for further analysis.

The volcano plot (Fig. 5) highlights the top 10 enriched and top 10 depleted taxa based on fold change (*p*-value < 0.05). The complete list of significantly enriched and depleted ASVs can be found in Supplementary Tables 3 and 4.

**Figure 5.**
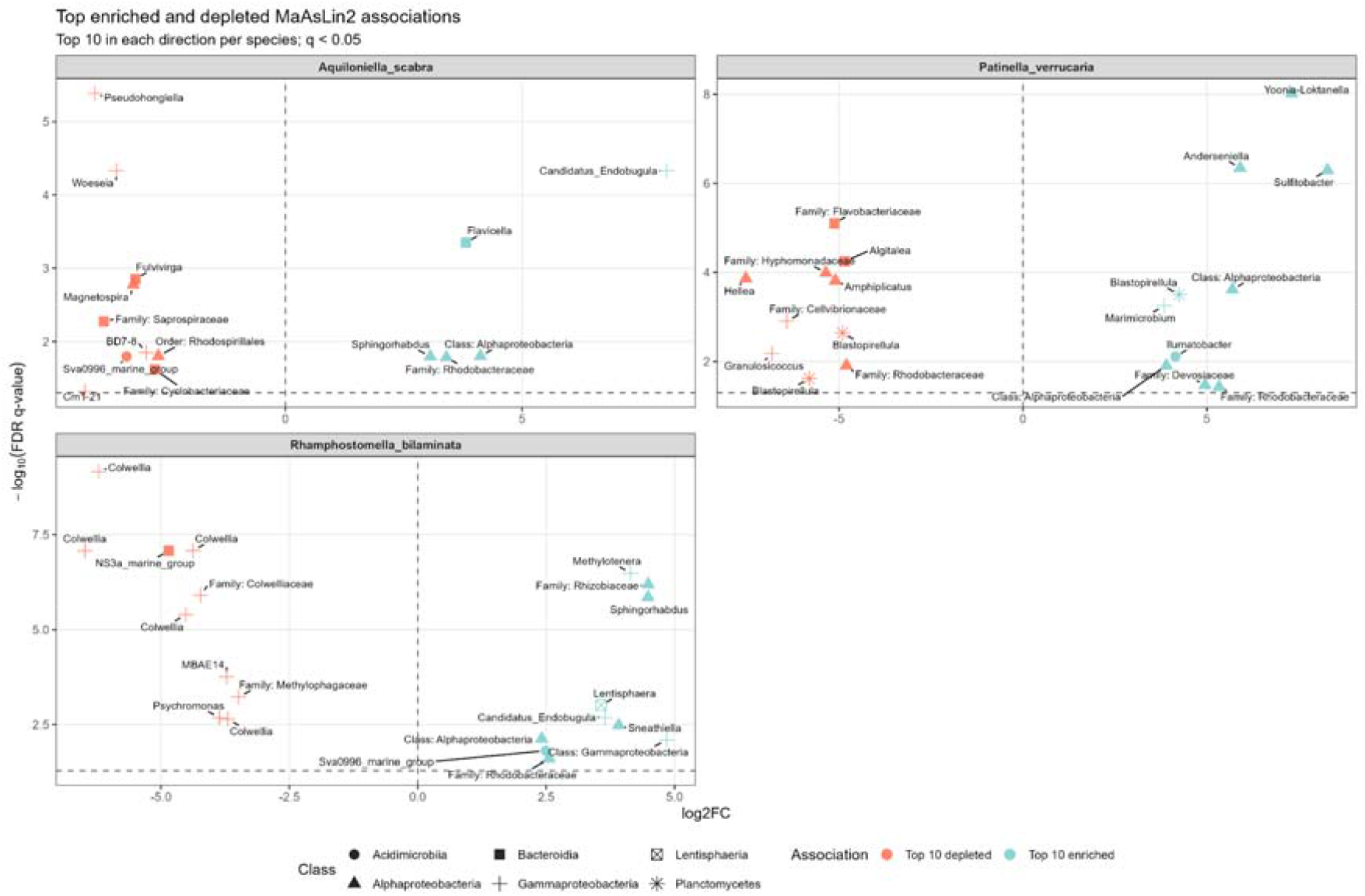
Volcano plots with top ten enriched and depleted taxa for each host species.

The “*Ca*. Endobugula” genus is strongly enriched in *A. scabra* (log_2_FC = +8.056) and in *R. bilaminata* (log_2_FC = +3.645) but is absent in *P. verrucaria* (Table 2). *Alphaproteobacteria* (*Rhodobacteraceae, Rhizobiales, Sphingomonadales*) are consistently enriched across all hosts, but *P. verrucaria* shows the highest diversity of enriched genera within this class (e.g., *Sulfitobacter, Yoonia⍰Loktanella, Anderseniella*).

**Table 2.**
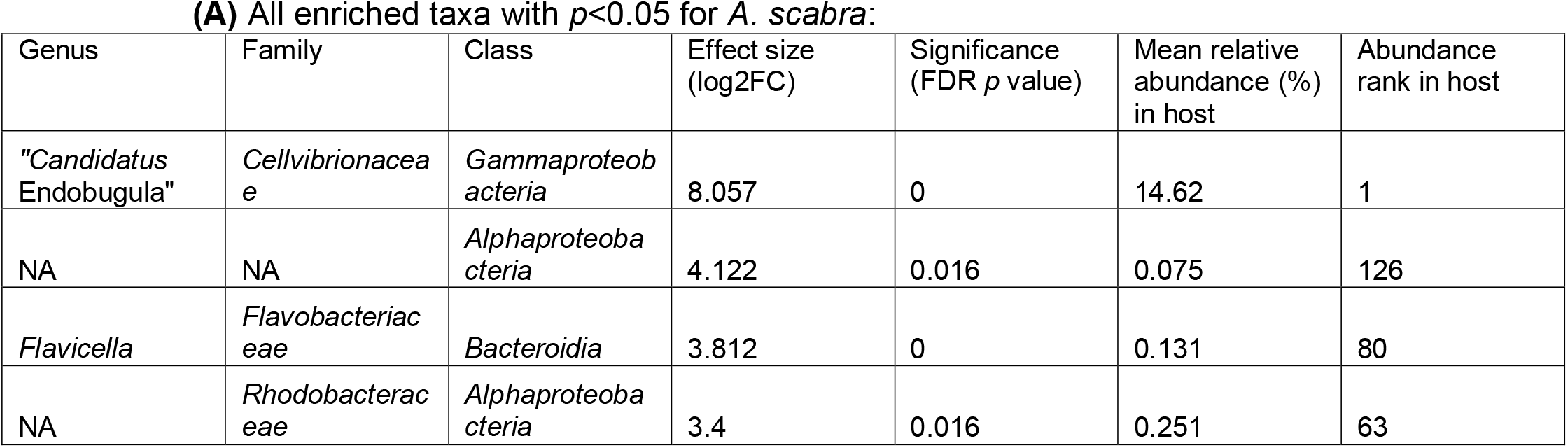
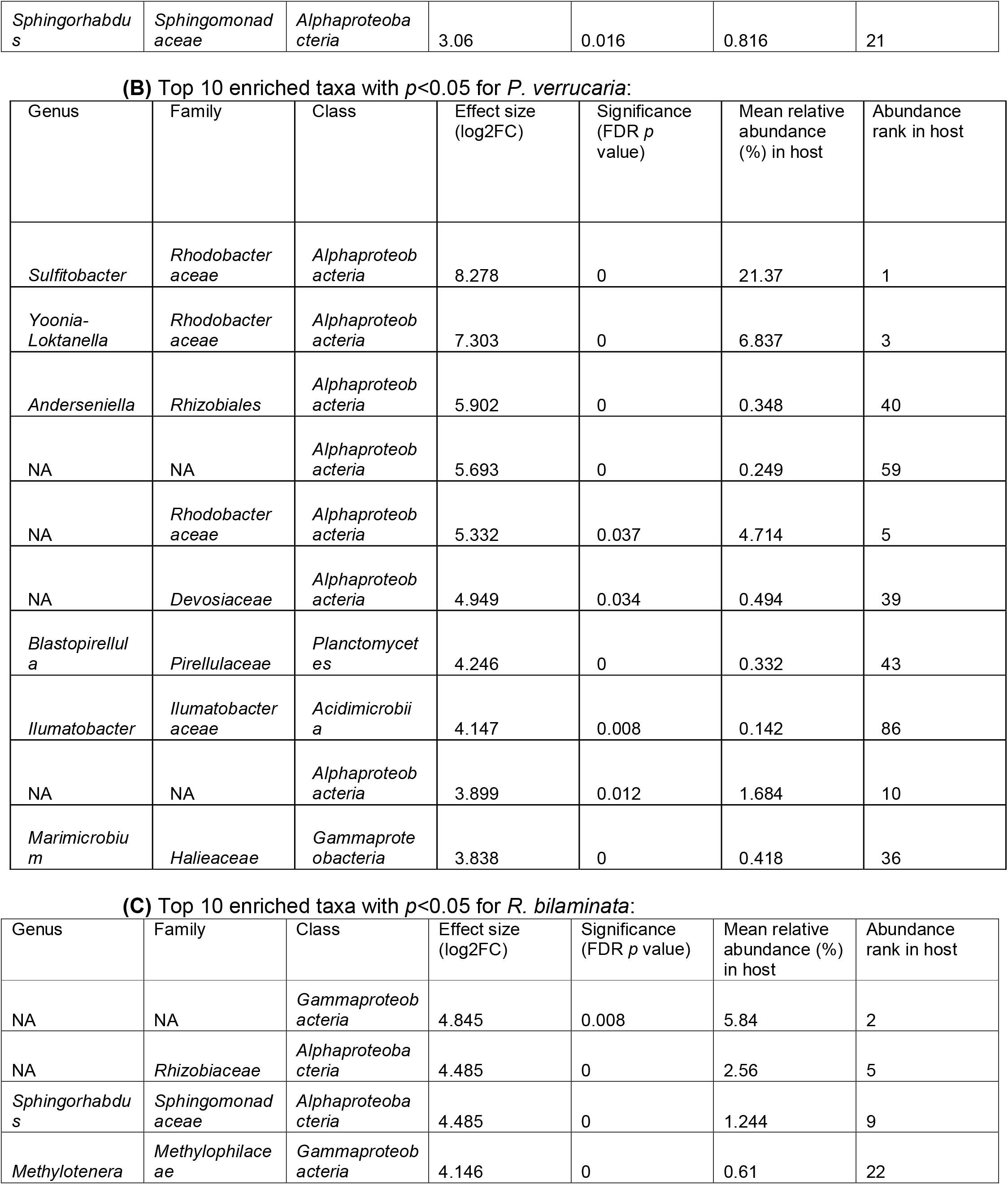
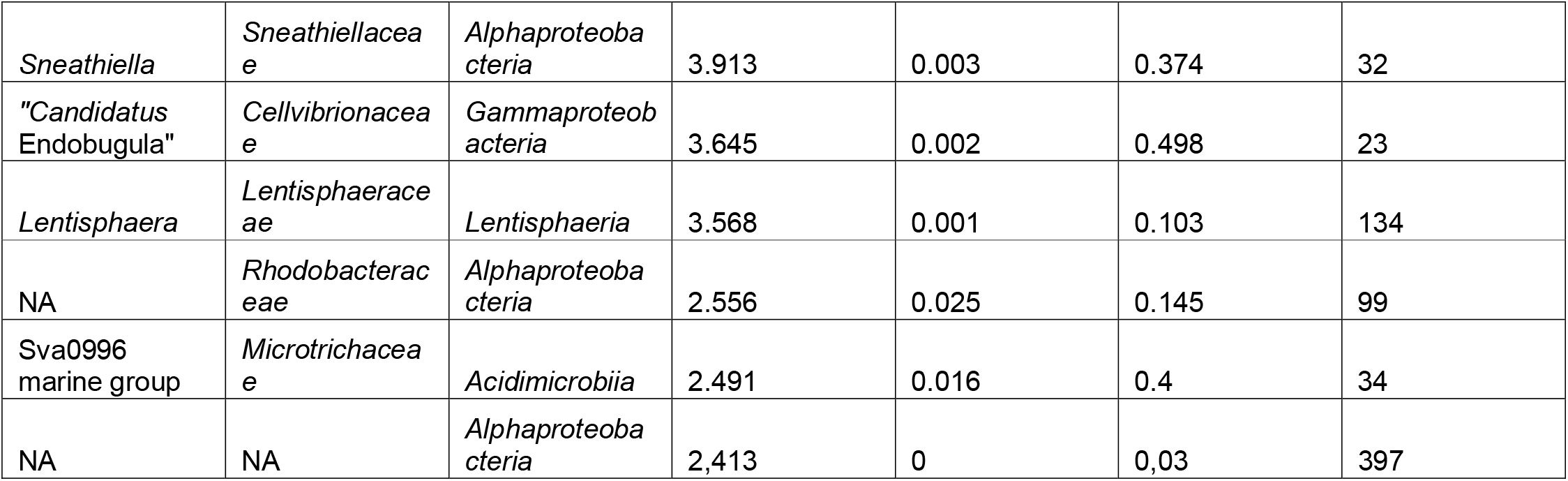
Key enriched taxa for three host species. NA – ASV that could not be assigned to a genus or family.

A notable feature across all differentially abundant taxa is the large number of ASVs that could not be assigned to a genus (marked as NA or assigned only at higher ranks such as family, order, or class). This is particularly evident in the enriched taxa of *R. bilaminata*, where several top hits are identified only at the class level. This suggests that a substantial fraction of the bryozoan-associated microbiota remains uncharacterized, likely representing novel or understudied bacterial lineages that may be specific to these hosts and merit further investigation.

To examine how the level of enrichment varies with microbial prevalence, we plotted log2FC against the abundance rank for each host species (Fig. 6)

**Figure 6.**
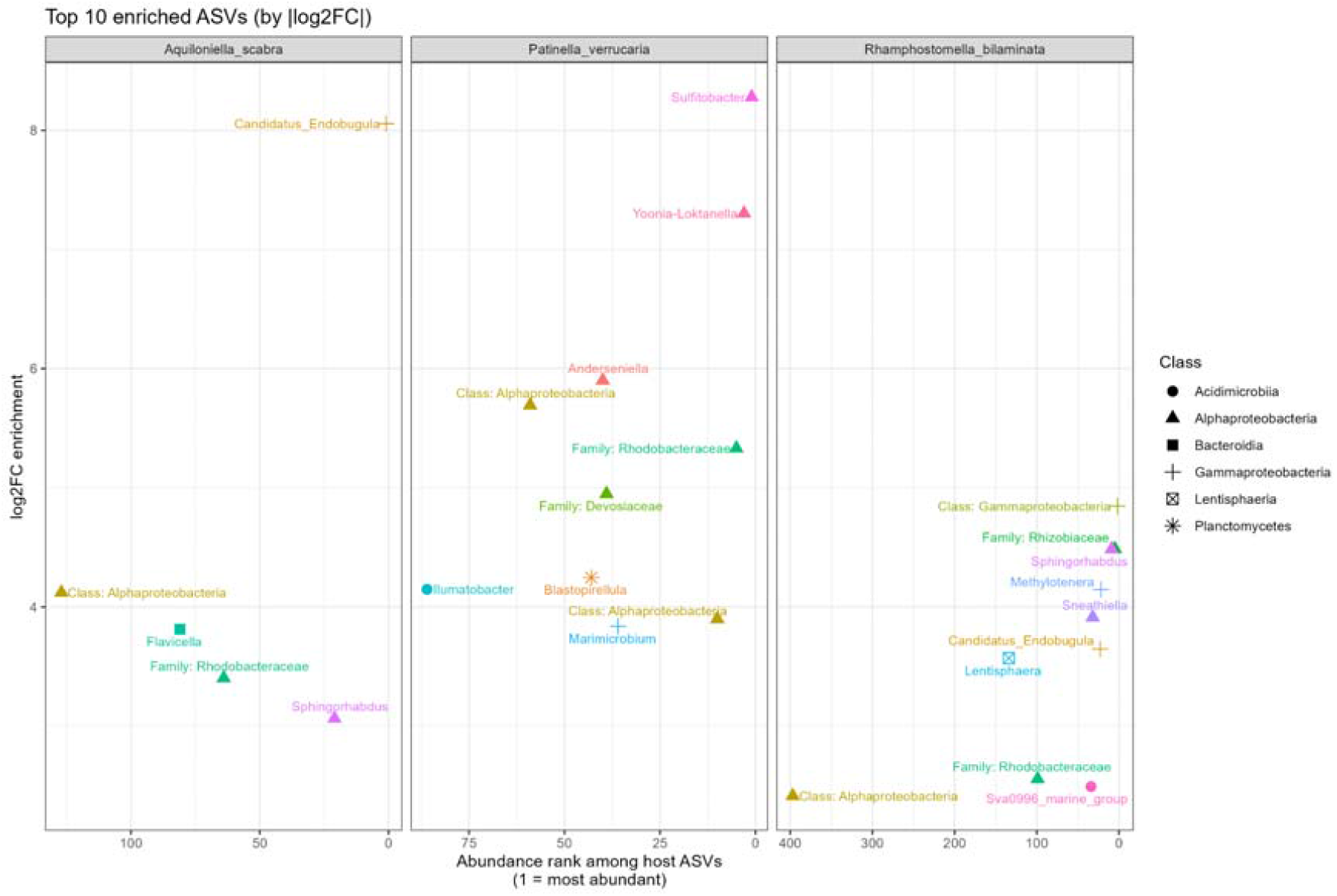
Key enriched taxa sorted by the abundance rank.

## Discussion

### Overall trends

*A. scabra* hosts only one strongly enriched and highly abundant bacterium, “*Ca*. Endobugula”, and several enriched but low abundance species. *R. bilaminata* hosts a community without a single dominating microorganism but also features the largest number of depleted ASVs. *P. verrucaria* has the largest number of enriched taxa, and in particular, three strongly enriched and highly abundant bacteria from the family *Rhodobacteraceae* (Supporting data of [32]): *Sulfitobacter, Yoonia-Loktanella*, and one not assigned to a genus.

### Enriched taxa

Majority of the significantly enriched bacterial taxa are known to be associated with marine invertebrates, supporting possible symbiotic relationship rather than incidental co-occurrence. Most important among them is “*Ca*. Endobugula”, detected in two hosts (*A. scabra* and *R. bilaminata)*, uncultivated genus of *Gammaproteobacteria* that is a well-established symbiont of bryozoans [52] and is best known as the source of the bryostatins, complex polyketides, assembled by a modular polyketide synthase (PKS) gene cluster, that act as potent protein kinase C (PKC) modulators with anticancer and neurological activity [53].

Members of the family *Rhodobacteraceae* were recovered across all three hosts and included *Yoonia*-*Loktanella, Sulfitobacter* (in *P. verrucaria*), and several unidentified genera (in all three hosts). This family is a recurrent symbiont of corals [54], sponges [55], and brittle stars [56], and its members have been shown to encode polyketide biosynthesis pathways in sponges [55]. *Rhodobacteraceae* representatives were also previously identified in all three stages of the *P. verrucaria* life cycle, indicating vertical transmission of the symbiont [32]. Moreover, this family was the second most abundant family detected in another Arctic bryozoan, *Tricellaria ternata*, and included genera *Yoonia*-*Loktanella* and *Sulfitobacter* [30].

Consistent with this pattern, *Anderseniella* (in *P. verrucaria*) is a sponge symbiont whose genome is predicted to encode secondary-metabolite (polyketide) biosynthesis, and it was found within the same community as various *Rhodobacteraceae*, among them the genus *Loktanella* [55]. *Sphingorhabdus* (in all three hosts) likewise is associated with marine hosts such as sponges [55] and gorgonian corals [57]. *Methylotenera* (in *R. bilaminata*), *Ilumatobacter* (in *P. verrucaria*), *Blastopirellula* (in *P. verrucaria* and *R. bilaminata*) and Sva0996 marine group (in *P. verrucaria* and *R. bilaminata*) were detected in the bryozoan *T. ternata* [30]. *Ilumatobacter* was also reported as a symbiont in crab gills [58] and *Methylotenera* was found in marine copepods [59]. Sva0996 marine group has been poorly explored; however, it was previously detected in sponges [60]. Representatives of *Sneathiella* (in *R. bilaminata*) were isolated from nudibranchs [61] and sea anemones [62]. *Marimicrobium* (in *P. verrucaria*) and *Sphingorhabdus* (in all three hosts) were detected in an Antarctic sponge along with *Blastopirellula, Ilumatobacter, Anderseniella* and Sva0996 [63].

Some of the enriched taxa may reflect the environmental pool rather than host selection, with two distinct acquisition routes. The first one is the diet. Bryozoans are suspension feeders that capture particles from the water column, so a relevant source might be phytoplankton and its attached bacteria. *Sulfitobacter, Yoonia* and *Sphingorhabdus* are co-isolated members of the diatom phycosphere, and strains of all three stimulate growth of the diatom partner [64]. *Rhodobacteraceae* are early and persistent colonizers during assembly of a diatom microbiome [65].

The second route is contact with surfaces and suspended detritus, and it must be invoked for the macroalgal associations. Planctomycetes form a biofilm on the surface of macroalgae by attaching to it and utilizing the sulfated polysaccharides of the algae under the action of sulfatase [66]. *Devosia rhodophyticola* and *D. algicola* were isolated from a marine red alga [67]. *Anderseniella* is additionally an indicator genus of the endophytic bacterial community of the brown macroalga *Sargassum thunbergii*, and Sva0996 was also amongst the most dominant genera in both the algal body and the receptacles [68]. *Ilumatobacter* is one of only seven amplicon sequence variants universally present in the core microbiome of three intertidal *Fucus* congeners, alongside a *Blastopirellula* [69].

Macroalgal detritus is a quantitatively important particulate food source for coastal suspension feeders, contributing an estimated 59% of the diet of blue mussels at one Irish site [70], so degrading algal particles and their attached bacteria do enter the suspended pool. Bryozoans are also common epibionts of macroalgae, dominating kelp epibiont assemblages together with crustaceans, molluscs, cnidarians, and annelids [71, 72]. Two mechanisms — filter feeding with microalgae and contact macroalgae detritus — can deliver these bacteria to a bryozoan.

Overall, environmental acquisition is the normal establishment route for many well-characterized marine symbioses, and it is compatible with specificity of a particular symbiosis. While “*Ca*. Endobugula” is presumably a vertically maintained bryozoan symbiont [22] (but criticized in [25]), the remaining genera with invertebrate records, namely, members of the *Rhodobacteriaceae* family, *Anderseniella, Sphingorhabdus, Ilumatobacter* and *Sneathiella*, are known from other invertebrate hosts but are equally prominent on kelps, so isolation from an animal sample is not necessarily an evidence of host selection. The observed genera without invertebrate records are explained by the same algal pools. The *Roseobacter* clade, which includes both *Sulfitobacter* and *Yoonia-Loktanella*, is a partial exception: in brittle stars it has been detected in the cuticle rather than the gut, and it produces polyketides that may benefit the host [56], arguing for a closer association than simply diet. In the bryozoan *P. verrucaria*, bacteria of the *Roseobacter* clade were found in colonies (in the amoebocytes) and tissues (including placental), what would ensure direct transmission of the symbionts to the incubated larvae [32].

Interestingly, genera significantly enriched in *P. verrucaria* (*Yoonia-Loktaniella, Sulfitobacter, Ilumatobacter, Blastopirellula, Rubritalea*, Sva0996 marine group) and in *R. bilaminata* (*Methylotenera, Blastopirellula*, Sva0996 marine group) were also found to be abundant in the bryozoan *T. ternata* that is confamilial with *A. scabra* [30].

*Sulfitobacter* and *Yoonia-Loktaniella*, two genera dominant in *P. verrucaria* by the logFC and relative abundance percentage (21% and 7% of relative abundance, respectively), were present in *T. ternata* in much smaller quantity (1.7% and 2.2% of relative abundance, respectively). *Ilumatobacter, Blastopirellula, Rubritalea*, Sva0996 marine group and *Methylotenera* were all present in *T. ternata* in larger quantities than in *P. verrucaria* or *R. bilaminata*. The absence of a dominating symbiont and the equal distribution of present genera suggest weaker selection in the *T. ternata* host. Similarly, our microscopic studies were not able to detect bacterial bodies in congeneric *T. gracilis* and *T. arctica* (unpublished data), that demonstrates a mosaic distribution of the symbiotic associations, especially given that *Tricellaria* is closely related to *Aquilloniella* [73].

### Novel “*Ca*. Endobugula” representatives

*“Ca*. Endobugula” was identified in *A. scabra* (ASV_ASca) with the mean relative abundance of 14.6% across all host replicates) and *R. bilaminata* (ASV_RBil with the mean abundance of 0.5% across all host replicates). Both ASVs, along with nine additional ASVs affiliated with this genus, were also detected in the environmental controls, but at substantially lower mean relative abundances (Supplementary Table 1). Nine additional ASVs occurred only sporadically, appearing in a subset of replicates from a single habitat.

Previously, “*Ca*. Endobugula sertula” was identified in the sea water collected near *B. neritina* colonies [20]. The consistent presence of *“Ca*. Endobugula sertula” in the sea water collections supports the argument that it can be horizontally transmitted between the colonies of *B. neritina*. This argument is further supported by latitudinal variation in the presence of the symbionts in *B. neritina* colonies: it is assumed that “*Ca*. Endobugula sertula” has been re-acquired recently in one of the studied populations [74]. Also, the symbiont’s genome shows no reduction — such as small size, abundant pseudogenes, or low coding density — which is consistent with at least a periodical free-living state [24].

Within the *B. neritina* species complex, the association is patchy: several sibling species harbor “*Ca*. Endobugula” while others do not, indicating that the symbiosis is dynamic across hosts [74–76]. At that, the symbiont’s distribution has a biogeographic structure: it is more often present in lower-latitude *B. neritina* populations and tends to be absent at higher latitudes, a gradient interpreted as a stronger predation pressure closer to the equator, maintaining selection for the defensive mutualism [74, 75]. Our results contradict this theory: the White Sea is a high-latitude, subarctic system, and based on the above reasoning the defensive “*Ca*. Endobugula” symbiosis should have been rare or absent, but it is not. They also suggest that the protective role of “*Ca*. Endobugula” could be much more extensive than expected. To prove this, larvae of *A. scabra* and *R. bilaminata* should be further studied by molecular methods.

## Conclusions

To date, “*Ca*. Endobugula” symbionts have been characterized exclusively from two species of the genera *Bugula* and *Bugulina* (Bugulidae). Our study identifies two new “*Ca*. Endobugula” representatives in two new bryozoan hosts, *A. scabra* and *R. bilaminata*. Sequencing of the genomes of these symbionts may yield insights into the biochemical, genetic, ecological, and evolutionary aspects of bryostatin synthesis.

Further, we identified *Rhodobacteriaceae* representatives that are enriched in all three studied bryozoan hosts. In *P. verrucaria*, we identified several *Rhodobacteriaceae* representatives, including *Sulfitobacter* and *Yoonia-Loktanella*, that are known for producing polyketides in sponges. However, a substantial proportion of the *Rhodobacteriaceae* representatives could not be classified at the genus level, likely reflecting the limited taxonomic characterization of this group in marine invertebrates.

Overall, the bryozoan microbiome appears to be assembled from two environmental sources: bacteria associated with algae, likely acquired through the attachment substrate and diet, and bacteria taken up from the surrounding seawater. Against largely environmentally acquired background in bryozoan hosts, some bacteria are suggested to be capable of vertical inheritance, including “*Ca*.Endobugula”. Yet the resulting community is not a passive reflection of the host surroundings since it differs significantly in composition from the environmental communities from which it is drawn.

## Acknowledgements

We gratefully acknowledge Alla Shevchenko for the contribution in DNA extraction and library preparation.

Sampling was carried out using the resources of the Educational and Research Station “Belomorskaia”, and microscopic examinations were performed using the facilities and equipment of the Department of Invertebrate Zoology and the Resource Centre “Development of Molecular and Cell Technologies” at the Saint Petersburg State University. The authors are grateful to Serge Bagrov, Saint Petersburg State University, for his help with sampling.

Financial support was provided by the Russian Science Foundation (grant №23-14-00351-P) (sequencing and data analysis). Additional data analysis and final preparation of the manuscript were supported by the Saint-Petersburg State University (research project 153213231). Publication has been supported in part by the Russian Science Foundation (grant №24-14-00276).

## Author contributions

AD – Formal analysis (16S rRNA gene amplicon data processing and statistical analysis), Visualization, Data curation, Conceptualization, Writing – original draft

ET – Investigation (sample processing, DNA extraction, PCR, and library preparation)

EB – Investigation (sample processing, electron microscopy), Visualization (electron microscopy), Conceptualization

OK – Investigation (sample collection and processing), Visualization (light microscopy), Writing – sampling sites and sample collection procedures

AV – Investigation (sample processing, electron microscopy), Visualization (electron microscopy)

MG – Supervision, Data curation, Writing – review and editing

AO – Supervision, Conceptualization, Writing – review and editing

All authors have read and approved the final version of the manuscript.

## Data availability

Raw sequencing reads are deposited to the GenBank SRA database under the Bioproject number PRJNA1519290. Processed data and code are available in the GitHub repository: https://github.com/annadukat/Microbiomes-of-the-Bryozoans-from-the-White-Sea

