## Supplementary Figures for "Novel “*Candidatus* Endobugula” hosts and distinct microbiomes in the bryozoans from the White Sea"

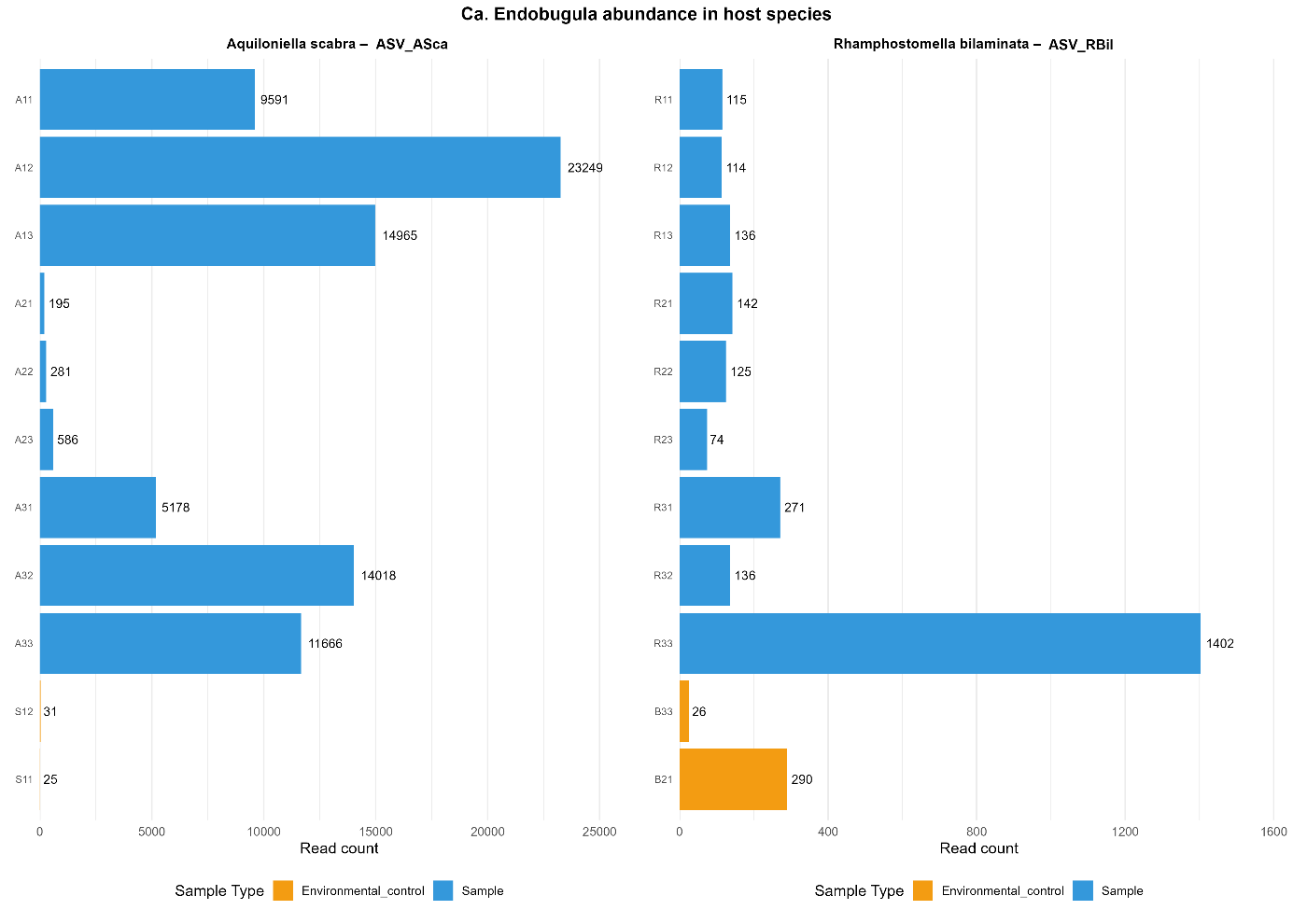

**Supplementary Figure 1.** "*Ca.* Endobugula" abundance in the host species. Y-axis represents sample id, X-axis represents read counts.

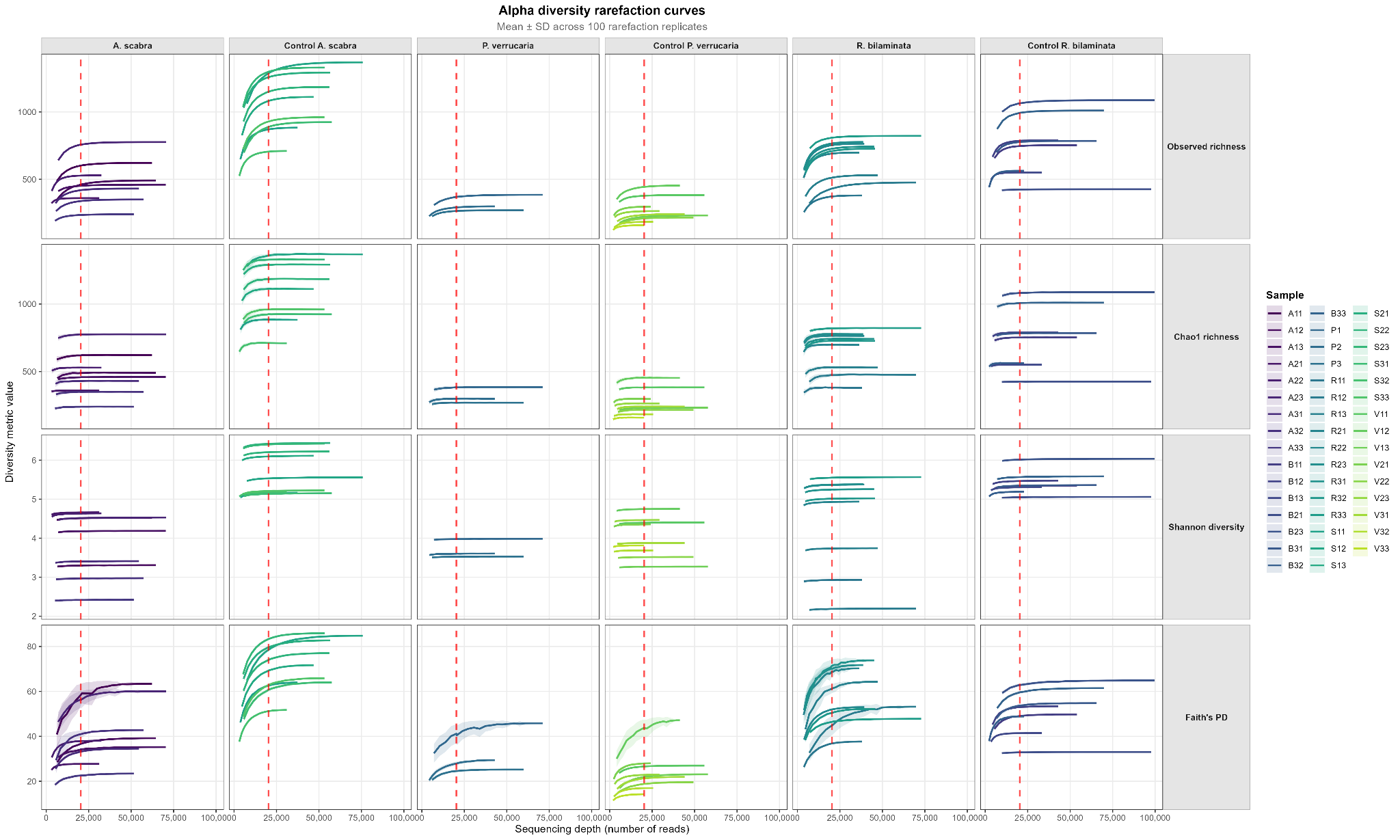

**Supplementary Figure 2.** Rarefaction curves showing the alpha-diversity indices across all samples. The dashed line indicates the minimal library size among all samples.

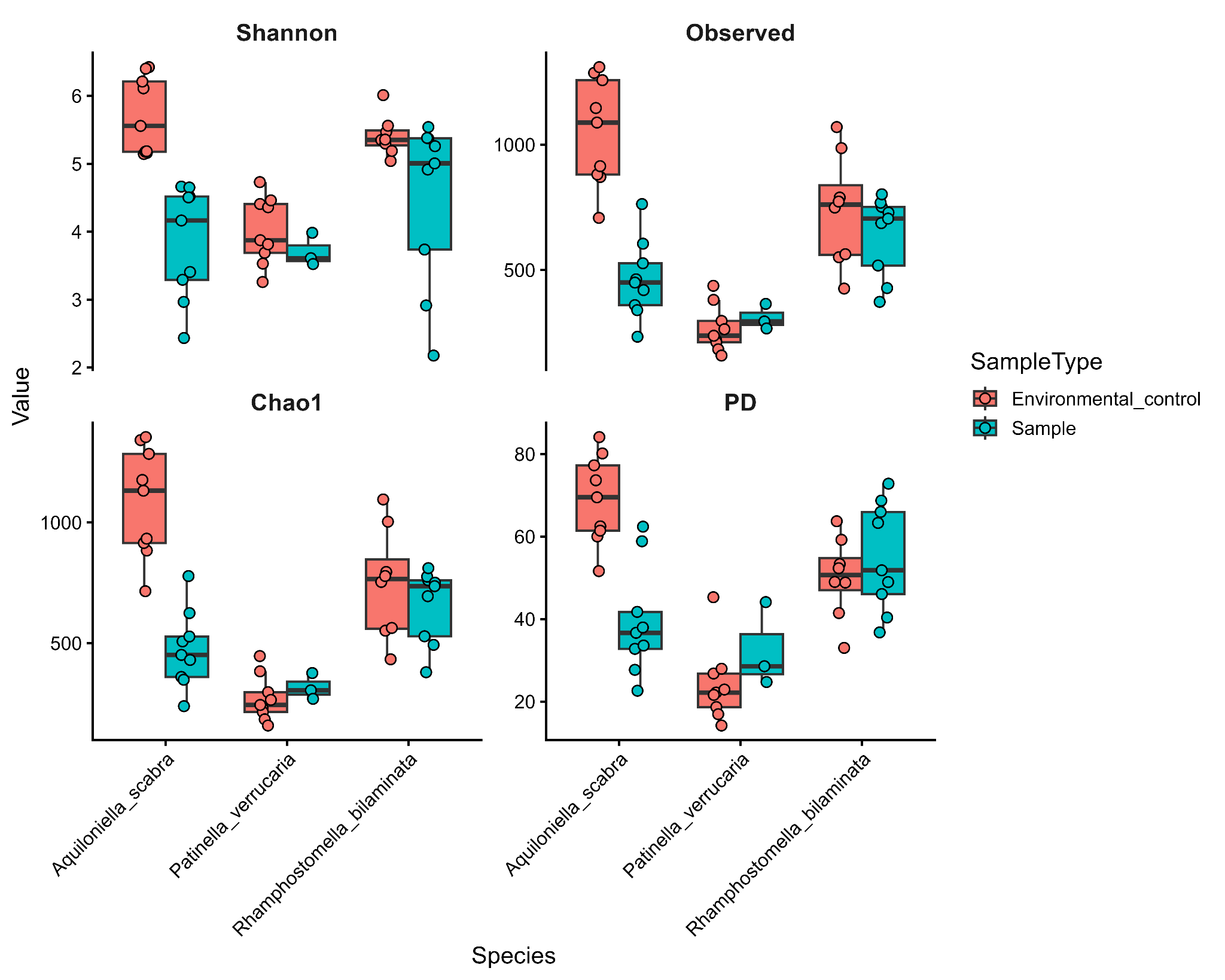

**Supplementary Figure 3.** Alpha-diversity box plots of bryozoan samples compared with respective environmental controls. The normalized library size was set to the minimum library size.
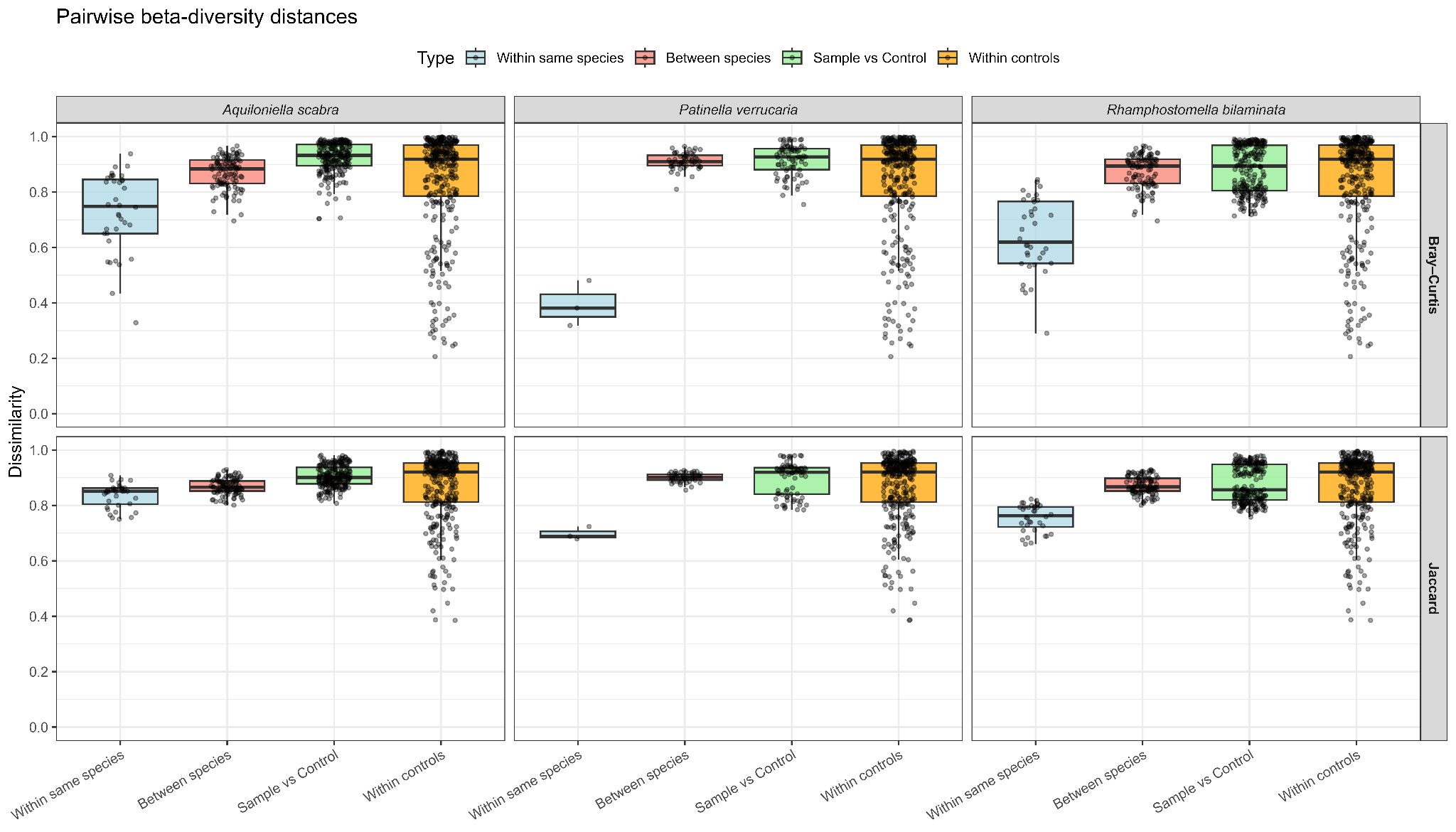
 **Supplementary Figure 4.** Pairwise beta-diversity distances for all species. “Within species” represents pairwise comparisons among samples from the same host species; “Between species” represents comparisons between samples from one bryozoan species and samples from the other two bryozoan species; “Sample vs Control” represents comparisons between samples from each host species and their respective controls; and “Within controls” represents pairwise comparisons among all control samples.
